# Thermal compensation reduces DNA damage in UV-exposed amphibian larvae: Implications for high latitudinal and altitudinal species

**DOI:** 10.1101/2023.06.25.546466

**Authors:** Coen Hird, Rebecca L. Cramp, Craig E. Franklin

## Abstract

1. Increases in ultraviolet radiation (UVR) correlate spatially and temporally with global amphibian population declines and interact with other stressors such as disease and temperature. Declines have largely occurred in high-altitude areas associated with greater UVR and cooler temperatures.

2. UVR is a powerful mutagenic harming organisms largely by damaging DNA. When acutely exposed to UVR at cool temperatures, amphibian larvae have increased levels of DNA damage. Amphibians may be able to compensate for the depressive effects of temperature on DNA damage through thermal acclimatisation, but it is unknown whether they or other ectotherms have this capacity.

3. We reared striped marsh frog larvae (*Limnodynastes peronii*) in warm (25°C) and cool (15°C) temperatures under either a low or moderate daily dose of UVR (10 and 40 µW cm^-2^ UV-B for 1 h at midday, respectively) for 18-20 days and then measured immediate DNA damage resulting from an acute high UVR dose (80 µW cm^-2^ UV-B for 1.5 h) at a range of test temperatures (10, 15, 20, 25, and 30°C).

4. Larvae acclimated to 15°C and exposed to UVR at 15°C completely compensated UVR-induced DNA damage compared with 25°C acclimated larvae exposed to UVR at 25°C. Additionally, warm-acclimated larvae had higher CPDs than cold-acclimated larvae across test temperatures, which indicated a cost of living in warmer temperatures. In contrast, larvae reared under chronic elevated UVR levels showed no evidence of UVR acclimation resulting in lower DNA damage following an acute high UVR exposure.

5. Our finding that thermal acclimation in *L. peronii* larvae compensated UVR-induced DNA damage at low temperatures suggested that aquatic ectotherms living in cool temperatures may be more resilient to high UVR than previously realised.

6. We suggested individuals or species with less capacity for thermal acclimation of DNA repair mechanisms may be more at risk if exposed to changing thermal and UVR exposure regimes but cautioned that thermal acclimation of DNA repair mechanisms may not always be beneficial.

## INTRODUCTION

Changes in stratospheric ozone over the past four decades due to anthropogenic emissions have altered solar ultraviolet radiation (UVR) conditions at Earth’s surface, interacting with other aspects of climate change in complex ways to affect human health, food and water security, organisms, and ecosystems (Barnes et al., 2022). Early models forecasted significant increases in UVR reaching Earth’s surface across all latitudes, leading to a surge in scientific research surrounding UVR effects on biota (Barnes et al., 2019). Many of the predicted adverse effects of changing UVR have been avoided because of the Montreal Protocol with its amendments and adjustments that halted the production of ozone-depleting substances (Morgenstern et al., 2008; World Meteorological Organization, 2022). In spite of these interventions, ozone depletion has led to a 2-5% increase in UVR levels since 1950 (Lemus-Deschamps & Makin, 2012; Madronich et al., 1995) and continues to drive climate change in the Southern hemisphere (Arias et al., 2021; Bais et al., 2015; Barnes et al., 2022; Blunden & Boyer, 2021; Fahey et al., 2018; Meinshausen et al., 2020; Montzka et al., 2018; Williamson et al., 2014). Critically, climate change has become a more important determinant of changing surface UVR levels than ozone depletion (Bais et al., 2018; Barnes et al., 2022). The increasing frequency of extreme climate events such as heatwaves, fires, droughts, floods, storms, and rain (Knuesting et al., 2018; Rammer et al., 2021; Watermann et al., 2020), changing snow/ice (Cabrera et al., 2012) and phenology (Cohen et al., 2018) can all serve to increase UVR levels reaching organisms and ecosystems.

In aquatic ecosystems, climate change-associated increases in solar UVR levels are particularly concerning for ectothermic animals (Paaijmans et al., 2013; Sokolova & Lannig, 2008). Most groups of aquatic ectotherms show susceptibility to damage by UVR at the cellular level, leading to population level impacts (Peng et al., 2017). Solar UVR has wide-ranging effects on organisms and biological processes. UVR can penetrate significantly into aquatic systems, particularly those with clear water and little overhanging vegetation (Xenopoulos & Schindler, 2001) at high altitudes and latitudes (Pfeifer et al., 2006; Wang et al., 2014). UVR is a mutagenic and cytotoxic agent that can directly damage nucleic acids and cause oxidative insult to lipids and proteins (Cadet & Richard Wagner, 2013; Lesser, 2006). Exposure to UVR results in the formation of pyrimidine (6-4) pyrimidone and cyclobutene pyrimidine dimers (CPDs) between nucleotide bases that can disrupt gene expression patterns and lead to mutations (Goodsell, 2001; Tevini, 1993). If DNA damage occurs at rates that exceed DNA repair rates, deleterious photoproducts may accumulate, resulting in whole-organism fitness consequences (Pandelova et al., 2006; Schuch et al., 2015). UVR-associated DNA damage can be repaired via nucleotide excision repair (NER) or photoenzymatic repair (PER) in some organisms (Davies, 1995).

Increases in the amount of UVR reaching the Earth’s surface and global warming correlate spatially and temporally with many historic amphibian population declines (Berger et al., 1998; Carey, 1993; Lips et al., 2006). Over the past few decades amphibian extinction rates exceeded, and are predicted to continue to exceed, background rates by over four orders of magnitude (Alroy, 2015). Although habitat loss/fragmentation and over-exploitation remain the primary threats to global amphibian populations, a significant number of population declines have occurred in pristine montane environments where temperatures are lower and UVR levels higher (Biodiversity Group, 1999; Blaustein & Wake, 1990, 1995; Bradford, 1991; Kiesecker et al., 2001; McDonald, 1990; Richards et al., 1993; Young et al., 2001). Most of these declines have been linked to the emergence and spread of two novel pathogenic amphibian chytrid fungi *Batrachochytrium dendrobatidis* (Bd) and *Batrachochytrium salamandrivorans* (BSal) (Berger et al., 1998; Gillespie et al., 2020; Martel et al., 2013; Young et al., 2001). However, amphibian declines are linked to multiple, often-interacting stressors, including UVR (Alton & Franklin, 2017; Bancroft et al., 2008; Blaustein & Kiesecker, 2002; Grant et al., 2016). Increased UVR was hypothesised to influence amphibian populations directly by having negative effects on embryos and larvae because these life stages are often diurnal and typically occur during spring and summer when UV-B levels are highest (Blaustein et al., 2003). Amphibians are particularly vulnerable to UVR exposure, showing increased mortality, morphological malformations in embryos, and slowed developmental and growth rates, physiological and behavioural changes, and reduced locomotor performance in larvae (see review by Alton & Franklin, 2017). The accumulation of UV-B-induced DNA damage is also a trigger for photo-immunosuppression (Kripke et al., 1992; Schwarz, 2005) and UVR is proposed to be immunosuppressive in amphibian larvae (Ceccato et al., 2016; Cramp & Franklin, 2018; Franco-Belussi et al., 2016, 2018; Londero et al., 2019). Consequently, factors that influence or limit the capacity of amphibians to repair UV-B-induced DNA damage, such as temperature, could have significant ramifications for host responses to pathogens such as *Bd*.

DNA repair rates are thermally sensitive (Lamare et al., 2006; Li et al., 2002; MacFadyen et al., 2004; Morison et al., 2019; Pakker, Martins, et al., 2000), and the negative effects of UV-B on amphibians are compounded when UV-B exposure occurs at low temperatures (Alton & Franklin, 2012; Bancroft et al., 2008; Kiesecker & Blaustein, 1995; Lundsgaard et al., 2020a; van Uitregt et al., 2007). Striped marsh frog (*Limnodynastes peronii*) larvae acutely exposed to UVR at cool temperatures showed increased DNA damage compared with larvae exposed to equal UVR levels at warmer temperatures (Morison et al., 2019), suggesting that the thermal sensitivity of PER (the primary mode of UV-B-associated DNA repair in amphibians; C. P. dos Santos et al., 2018; Mei & Dvornyk, 2015) may partially explain why a disproportionately high number of amphibian declines have occurred at higher altitudes (>400 m above sea level (ASL)) where UVR levels are higher and temperatures are cooler (Biodiversity Group, 1999; Blaustein & Wake, 1990, 1995; Bradford, 1991; Kiesecker et al., 2001; McDonald, 1990; Richards et al., 1993; Young et al., 2001). However, amphibian species differ in their thermal sensitivity of UVR-induced DNA damage accumulation (Hird et al., 2022). Critically, many organisms can acclimatise their thermal sensitivities in response to their environmental conditions through active phenotypic plasticity, remodelling their physiology to shift the thermal reaction norm for a given trait in order to compensate for detrimental thermal effects on physiological rates and organismal fitness (Angilletta, 2009; Rohr et al., 2018; Seebacher, White, et al., 2014). When predicting UVR risk in ectotherms from cool environs, or how ectotherms will respond when moving into environments with different thermal and UVR regimes, it is critical to consider whether these organisms have the capacity for acclimatisation to compensate for the depressive effects of temperature on DNA repair rates.

In this study, we examined the effect of rearing (acclimation) temperature (cool = 15°C, or warm = 25°C) on the thermal sensitivity of UVR-associated CPD accumulation in *L. peronii* larvae. In addition to their different thermal rearing environments, larvae were also concurrently exposed to either a low (10 µW cm^-2^) or moderate (40 µW cm^-2^) level of UV-B through the thermal acclimation period to assess the capacity for UVR hardening in the mitigation of DNA damage accumulation following high UVR exposure. It was hypothesised that larvae would demonstrate thermal acclimation of physiological function following chronic exposure to both low temperature and high UVR, ameliorating the depressive effects of low temperature on DNA damage accumulation and protecting against the negative impacts of increased UVR exposure.

## MATERIALS AND METHODS

### Experimental animals and general methods

Freshly laid *Limnodynastes peronii* spawn were collected near Meeanjin on Yuggera Country (−27.7856, 153.2702; QLD, Australia). Spawn was immediately transported to The University of Queensland, split into smaller clumps, and hatched in two-litre ice-cream containers half-filled with carbon-filtered Brisbane City tap water at either 15°C or 25°C. Water temperatures were achieved by holding spawn in separate temperature-controlled rooms where they were maintained for the duration of the experiment. Larvae were fed frozen spinach and partial water changes were conducted every second day. Larvae were reared under a 12L:12D photoperiod generated using standard fluorescent lights. Larvae held at 25°C hatched at two days post-collection, whereas larvae held at 15°C hatched at eight days post-collection. Upon hatching, approximately half of the larvae from each temperature treatment were assigned to either a low or moderate UV-B lighting treatment which corresponded to approximately 10 or 40 µW cm^-2^ at the water surface which was administered for 1h daily (at midday) using 40 W, full spectrum fluorescent lights which emit visible light, UV-A, and UV-B (Repti-Glo 10.0, 1200 mm, Exo Terra, Montreal, Canada). UVR for the experimental exposure was generated using the same lights. Light intensities for UVB were measured using a radiometer (IL1400BL, International Light Inc., Newburyport, USA; Table 1). UVA likely varied commensurate with UVB during the exposure period, but UVA was not directly measured. This is because CPDs are not generally attributed to UVA as it is only weakly absorbed by DNA (Rochette et al., 2003), and that we measured ‘light-CPDs’ which are predominantly generated by direct photodamage rather than UVA-induced reactive oxygen species (ROS) causing ‘dark CPDs’ (Cadet & Douki, 2018; Portillo-Esnaola et al., 2021; Premi et al., 2015).

**Table 1.** Measured UV-B values for each UVR chronic and acute treatment. Chronic (10 and 40 µW cm^-2^ treatments were exposed for 1 h daily. Acute (80 µW cm^-2^) UV-B exposure was for 1.5 h in Experiments 1 and 2, and for 1 h in Experiment 3. All values are reported in µW cm^-2^.

| Nominal daily UV-B dose | Nominal UV-B irradiance | Measured irradiance (mean ± SD) |
| --- | --- | --- |
| 36,000 | 10 | 9 ± 0.7 |
| 144,000 | 40 | 38.7 ± 1.2 |
| 432,000 | 80 | 73 ± 5.1 |

### Experimental design

Larvae were maintained under their treatment conditions for 18-20 days post-hatching since thermal acclimation typically occurs over several weeks (Bouchard & Guderley, 2003; Orczewska et al., 2010; Rogers et al., 2004). The experiment was repeated twice over consecutive days to accommodate the necessary technical and biological replication since space restrictions prevented all animals being exposed and treated simultaneously. Half of the larvae (n = 55) were individually placed into separate wells in six-well plates containing 10 ml of Brisbane filtered tap water. The plates were evenly distributed across five water baths (volume 32 L; L 650 mm x W 410 mm x H 210 mm; water depth ∼160 mm) at 10°C, 15°C, 20°C, 25°C, or 30°C (n = 22 larvae per test temperature: n = 10-12 per acclimation temperature and n = 5-6 per UVR acclimation treatment). The temperature of the experimental water baths was controlled using 300 W aquarium heaters (AquaOne) and water was circulated by small aquarium pumps to ensure thermal uniformity and to move plates across the surface of the water to homogenise the UV-B received by larvae. Larvae were left for one hour to adjust to the experimental temperature. UVR lights were switched on at 12:00pm and all larvae were exposed to ∼80 µW cm^-2^ UV-B for 1.5 hours (Table 1). Afterwards, larvae were euthanised in buffered MS222 (0.25 mg L^-1^; Ramlochansingh et al., 2014) and snap frozen at −80°C. The experiment was repeated the following day with another 55 larvae. Despite the age difference (as days post hatching) between larvae in the two temperature treatments, all larvae were in the same developmental stage (Gosner stage 25; Gosner, 1960) at the time of the final UVR exposure.

### CPD detection

Genomic DNA was extracted and purified from whole-larvae homogenates using a Qiagen DNeasy Blood and Tissue Kit (Qiagen Inc., Hilden, Germany) and quantified using a Qubit dsDNA High-Sensitivity Assay Kit (ThermoFisher Scientific Inc., Waltham, MA, USA). CPD concentrations were determined using an anti-CPD ELISA assay following the primary antibody manufacturer’s protocol (Mori et al., 1991; NM-ND-D001, clone TDM-2, Cosmo Bio Co., Ltd.). Briefly, tadpole DNA (0.4 ng µL^-1^) was loaded in triplicate into a protamine sulfate-coated 96-well plate and dried onto the plate overnight at 37°C. The following day, the plates were washed with 0.05% Tween-20 in phosphate buffered saline (PBS-T), blocked with 2% fetal bovine serum in phosphate buffered saline and then the primary anti-CPD monoclonal primary antibody was applied (1:1000 in PBS). The plates were then washed with PBS-T, and the secondary antibody applied (biotinylated goat anti-mouse IgG (QD209886, Life Technologies, USA). The signal from the secondary antibody was amplified using an HRP-conjugated streptavidin (1:10,000 in PBS; ab7403, Abcam, Cambridge, UK). Colour development was achieved with TMB substrate (1 mg L^-1^ in 0.05M phosphate-citrate buffer, pH 5.0) following Mori et al., 1991. Colour development was stopped after 30 minutes with 2M H_2_SO_4_ and absorbance determined at 450 nm in a DTX880 multimode detector (Beckman Coulter, MN, USA) using the SoftMax® Pro program (Version 7.1.0, Molecular Devices LLC, CA, USA). CPD concentrations were calculated from a standard dose-response curve of UVC-irradiated calf thymus (NM-MA-R010, Cosmo Bio Co., Ltd.) on each plate. CPD concentration is reported as units of UVCR-dose equivalents per 20 ng of DNA.

### Replication statement

| Scale of inference | Scale at which the factor of interest is applied | Number of replicates at the appropriate scale |
| --- | --- | --- |
| Individuals | Water baths | 10-11 |

### Analyses

All statistical analyses were conducted in the R statistical environment (R Core Team, 2021). Regression diagnostics were analysed using the performance package (Lüdecke et al., 2021). Models were two-tailed, assumed a Gaussian error structure and all models met the assumptions of the statistical tests used. α was 0.05 for all tests. The effect of wet body mass on CPD accumulation was accounted for by fitting wet body mass as a covariate in the model.

DNA damage thermal reaction norms were modelled using a second order polynomial regression with acclimation UV, acclimation temperature, and exposure temperature as fixed effects and mass as a covariate. There was no effect of UV acclimation levels on CPD concentration, so UV was dropped from the model and data points within each UV acclimation group were combined for the subsequent analyses. To explicitly test for thermal acclimation, data from larvae acutely exposed to 15- and 25°C were subset and analysed separately in a 2×2 factorial design using linear Type II sums of squares analysis with the car package (Fox & Weisberg, 2018). Post hoc comparisons were conducted using the emmeans package (Lenth, 2020). All estimates reported from the model are estimated marginal means adjusted for the effect of the covariate.

Q_10_ values were calculated from the 2×2 factorial design following a derivation of the Arrhenius equation (White et al., 2012). We assumed that high values of the response variable (CPD concentration) represented a worse fitness outcome for amphibian larvae, inverse to the typical pattern of T_OPT_ shifting left with cold acclimation i.e., Seebacher, White, et al., 2014). Because we expected to see an increase in temperature associated with a reduction in CPDs, we calculated Q_10(25-15)_ values rather than Q_10(15-25)_ values in order to preserve typical Q_10_ values for interpreting patterns of thermal acclimation (Huey & Berrigan, 1996).

## RESULTS

With exposure to UVR, CPD accumulation in *Limnodynastes peronii* larvae was highly sensitive to ambient temperature (figure 1; F_2,98_ = 27.65, *p* < 0.001) in both rearing temperature treatments (no interaction; F_2,98_ = 1.22, *p* = 0.3) with CPD concentration declining with increasing exposure temperature. Larval wet body mass of *L. peronii* was negatively correlated with the abundance of CPDs following acute UVR exposure (F_1,98_ = 6.13, *p* < 0.015). Overall, larvae chronically exposed to 15°C had significantly lower CPD abundance after 1.5 h of UVR exposure than larvae chronically exposed to 25°C (F_1,98_ = 29.06, *p* < 0.001).

**Figure 1.**
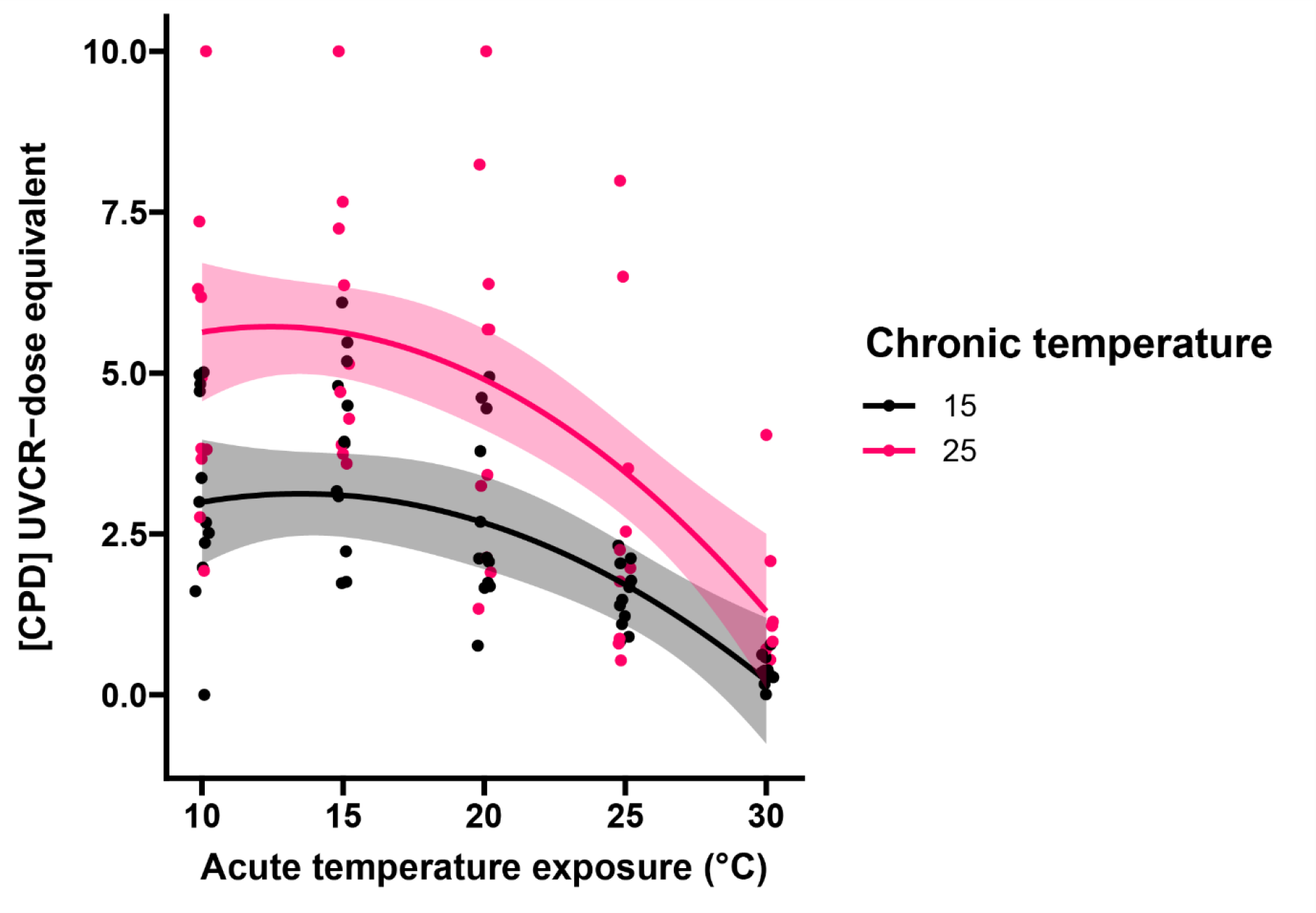
DNA damage (CPDs) immediately following 1.5 h 80 µW cm^-2^ UVB exposure in *Limnodynastes peronii* larvae reared at 15°C or 25°C and acutely exposed to a range of temperatures. Points are raw individual data points. Lines are mean predicted CPD concentration at a given test temperature for an average-sized *L. peronii* (8.5 mg). Shaded areas represent 95% confidence intervals for the fitted values.

Larvae reared at 15°C and acutely exposed to 25°C had lower levels of CPDs relative to larvae reared and tested at 15°C (Q_10(15-25)_ = 1.8). Conversely, larvae reared at 25°C and acutely exposed to UVR at 15°C had higher levels of CPD accumulation compared to larvae reared and tested at 25°C (Q_10(25-15)_ = 1.92). Larvae reared and tested at 15°C had significantly lower CPD levels compared with larvae acutely exposed to 15°C (F_2,98_ = 6.64, p = 0.014; figure 2) indicating that there had been beneficial thermal acclimation. There was no significant difference in CPD abundance between larvae exposed to UVR at their rearing temperatures indicating that full thermal compensation had occurred in larvae reared at 15°C.

**Figure 2.**
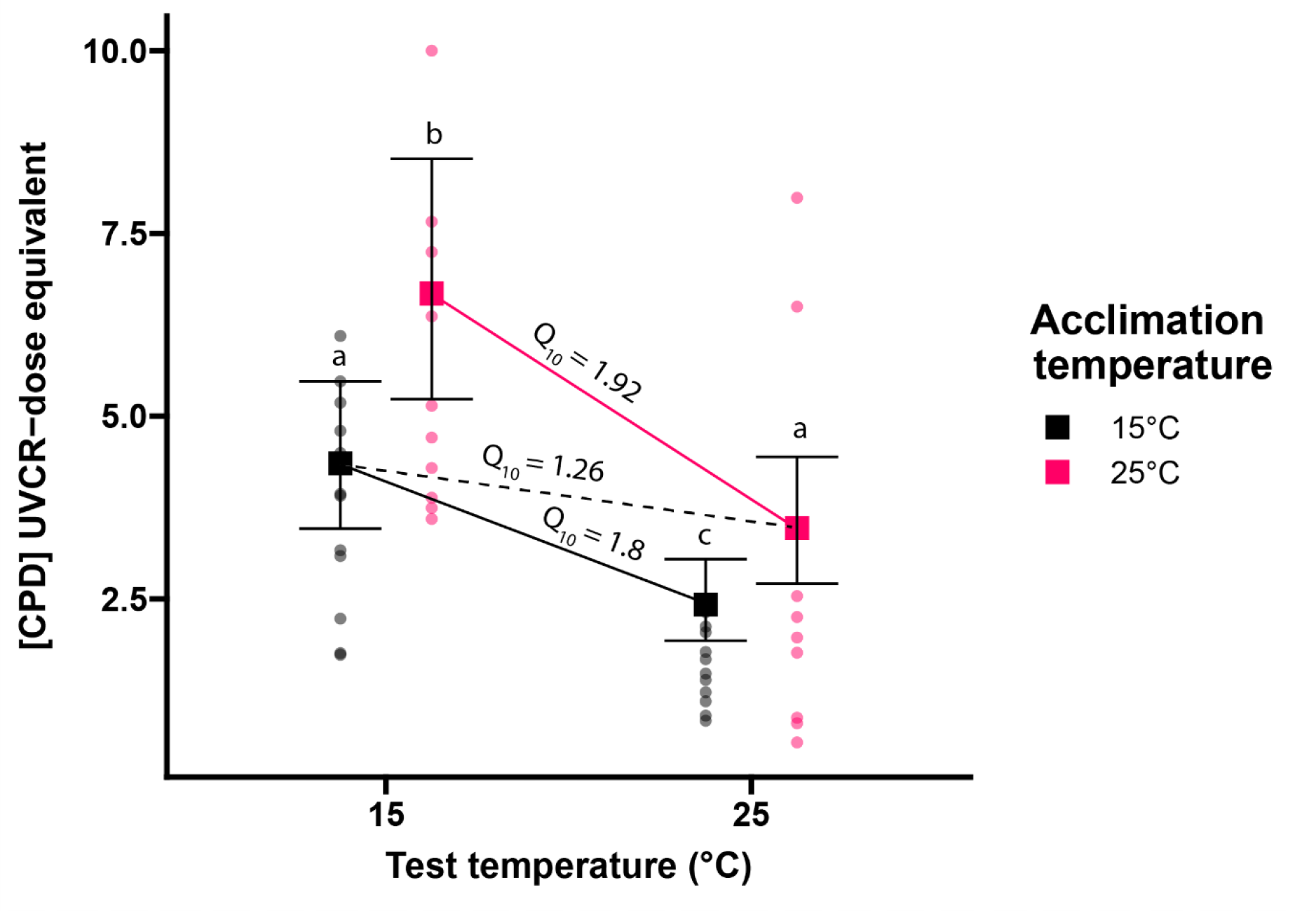
DNA damage (CPDs) immediately following 1.5 h 80 µW cm^-2^ UVB exposure in *Limnodynastes peronii* larvae reared at 15°C or 25°C and acutely exposed to either 15°C or 25°C. Round points are raw data points. Squares are mean predicted CPDs at a given test temperature for an average-sized *L. peronii* (8.22 mg). Error bars represent 95% confidence interval bounds for the for the fitted value. Letters represent significantly different groups. The solid black and red lines represent cold-acclimated and warm-acclimated larval acute thermal sensitivities, respectively. The dashed black line represents the post-acclimation thermal sensitivity.

## DISCUSSION

Our results demonstrated that CPD abundance following UVR exposure is highly thermally sensitive in *Limnodynastes peronii* larvae with a Q_10_ of ∼1.8 - 1.9. This is consistent with previous findings for this and other related amphibian species that show that UV-associated DNA damage repair rates are thermally sensitive (Hird et al., 2022; Morison et al., 2019). Importantly, larvae reared at low temperature were also able to offset the detrimental effects of protracted exposure to low temperatures through thermal acclimation of UV protection processes and/or DNA repair. To our knowledge, this is the first evidence of beneficial thermal acclimation of the processes associated with the management of DNA damage which allows animal to compensate for the depressive effects of low temperature on photolyase activity in ectothermic animals. This finding suggests that thermal compensation to low temperatures with acclimation can improve the tolerance of amphibian larvae to UVR. These data are supported in studies of plants, where thermal acclimation has been demonstrated to reverse the thermal dependence of UVR-induced DNA damage via photoreactivation (Pakker, Beekman, et al., 2000; Pakker, Martins, et al., 2000).

The finding that amphibian larvae can completely compensate UVR-induced DNA damage accumulation at low temperatures via thermal acclimation is particularly interesting considering that complete acclimation responses are relatively rare in ectotherms, and especially weak in amphibians (Havird et al. 2020). Complete compensation is also impressive considering that cold-acclimated animals were smaller than warm-acclimated animals in this study (mean mg ± S.D: 15°C acclimated = 7.6 ± 2.97; 25°C acclimated = 12.46 ± 7.73), because a given dose of UVR resulted in higher concentrations of whole body CPDs in smaller animals possibly due to a higher surface area to body volume ratio (Morison et al., 2020). In addition, acclimation responses in ectotherms tended to be immeasurable until over 30 days of acclimation (Havird et al., 2020), whereas the larvae in this study demonstrated strong acclimation responses after only 18-20 days. Because UVR is a potent selective pressure and that smaller larvae are more susceptible to the harmful effects of UVR (Morison et al., 2019), it is possible that cold-exposed larvae need to acclimate at a more rapid rate to avoid the deleterious effects of increased DNA damage occurring at low temperatures.

Across both thermal acclimation groups, CPD abundance was thermally sensitive at higher temperatures (> 20°C) and thermally insensitive at lower temperatures (< 20°C) suggesting that there may be an upper limit to the thermally sensitive accumulation of UV-associated CPD damage in *L. peronii* larvae. While our results show that acute exposure to warmer environments was associated with lower levels of UV-induced DNA damage overall, the finding that warm-acclimated animals had higher CPDs compared with cool-acclimated animals, irrespective of the exposure temperature, suggested that chronic exposure to lower temperatures provided some benefits to larvae that protected them against UV-associated DNA damage. Conversely, there may be a cost to animals from living at higher temperatures which increased their rate of DNA damage during UVR exposure. This apparent cross-tolerance may be important in natural systems, where climate warming might otherwise be expected to result in lower levels of UVR-associated DNA damage.

Thermal acclimation is not always beneficial. Mosquitofish with greater capacity for thermal acclimation to cool temperatures showed reduced swimming performance when in warm environments (Seebacher, Ducret, et al., 2014). Acclimation may also be associated with metabolic and other physiological trade-offs (Pörtner et al., 2006), such as the energetic costs involved in the synthesis of cutaneous melanin (Debecker et al., 2015). Trade-offs that occur in early life may have important repercussions for juvenile or adult amphibians after metamorphosis (Alton et al., 2012; Bancroft, 2007; Cramp et al., 2022; Lundsgaard et al., 2023; O’connor et al., 2014). Organisms can alter their phenology/behaviour in response to changing abiotic factors such as temperature (Chmura et al., 2019; Hill et al., 2021; Van Dyck et al., 2015; Woods et al., 2015). It is unknown whether thermal acclimation in aquatic ectotherms to minimise the effect of cold temperatures on UVR-induced DNA damage is associated with changes to behaviour and phenology or trade-offs with organismal fitness, though this warrants further investigation within and between species. If such a relationship exists, it could have serious implications for how freshwater ecosystems will respond to changing thermal and UVR environments in the future.

The physiological mechanisms of cold acclimation in *L. peronii* larvae that resulted in lower CPD abundance following UVR exposure remains unknown, though it may involve the upregulation of DNA repair genes, increased DNA repair enzyme activity, mobilising photoprotective agents such as melanin, or a combination of factors. Cold-acclimated larvae in this study were visibly darker than warm-acclimated larvae at the time of the acute high UVR exposure. Several studies show that UVA can indirectly cause ‘dark-CPDs’ through the formation of ROS (Cadet & Douki, 2018; Premi et al., 2015) from reaction with phaeomelanin, which may be thermally sensitive. However, eumelanin is the main pigment involved in the darkening response in amphibians (Frost-Mason & Mason, 2004; Giuseppe Prota, 1992), which exhibits a more photoprotective effect (Galván et al., 2011; Ito et al., 2018). Therefore, a darkening response during cold acclimation may reduce ROS-induced CPD formation. While higher temperatures are usually associated with acute increases in ROS production in ectotherms, some reptiles can acclimate to significantly reduce oxidative stress at high temperatures (Ritchie & Friesen, 2022). More research is needed to elucidate the relative contribution of light- and dark-CPDs following UVR exposure in amphibians from different thermal environments.

Unlike temperature, chronic exposure to a low daily dose of UVR (40 µw cm^-2^) did not lead to changes in CPD accumulation in *L. peronii* larvae following high intensity UVR exposure. This apparent lack of response is in contrast to the ‘UV hardening’ response that has been identified in plants, whereby the synthesis of compounds that absorb UVR (such as ‘sunscreens’, and flavonoids which also stimulate melanogenesis) are a primary acclimation response to UVR which provides protection against subsequent UVR exposure (Barnes et al., 2015; Jansen et al., 1998; Sen Mandi, 2016). Additionally, some plants exposed to UVR do not change photoprotective compound concentrations, but rather increase their capacity for DNA repair (Giordano et al., 2003). It is not well understood how, or even if, animals can acclimatise to UVR exposure. Our observation that *L. peronii* larvae were unable to acclimate to a moderate level of UV-B prior to a high intensity UV-B challenge may suggest a lack of UVR hardening capacity in high UVR environments. Alternatively, even low levels of UVR, such as those used in the low UVR treatment group, may have been enough to stimulate a UV hardening response with the higher UVR exposure not promoting any further response.

‘Enigmatic’ and disease-related amphibian declines typically occurred at high altitudes where UVR is greater, and temperatures are cooler. The compounding detrimental effects of cool temperatures and high UVR levels on amphibian health are well understood under acute exposure regimes (Lundsgaard et al., 2020b; Morison et al., 2019; van Uitregt et al., 2007). However, our data provides important evidence that amphibians living at high altitudes and in temperate environments could show plastic responses which compensate for the negative impacts of UVR and low temperatures on aquatic ectotherms. As climate change and ozone depletion continue to modify the thermal regimes and UVR exposures received by some UVR-sensitive freshwater ectotherms, capacity for thermal acclimation that reduces UVR-induced DNA damage may be evidence of resilience in the future (Seebacher, White, et al., 2014). Susceptibility to changing environmental conditions may be correlated to differential acclimation capacities. Species with greater plastic responses may be selected for in climates with increasing thermal fluctuations compared to species without a high degree of acclimation capacity.

## ACKNOWLEDGEMENTS

We acknowledge the Indigenous lands where the research from this article was conducted on, and realise sovereignty was never ceded. We acknowledge the cultural significance of amphibians in Indigenous cultures globally and recognise the relevance of amphibian conservation to Indigenous peoples. This research received support from an Australian Research Council Discovery grant awarded to CEF and RLC (DP190102152). CH was supported by an Australian Government Research Training Program Scholarship and a Ric Nattrass Research Grant from the Queensland Frog Society.

## DATA AVAILABILITY

The complete datasets and R scripts used for analysing the data are publicly available at UQ eSpace (doi.org/10.48610/221fbfe).

## AUTHOR CONTRIBUTIONS

Conceptualisation: RLC, CH, CEF; Methodology: CH, RLC, CEF; Validation: CH; Formal analysis: CH; Investigation: CH; Resources: CEF; Data curation: CH; Writing – original draft: CH; Writing – review & editing: CH, RLC, CEF; Visualisation: CH; Supervision: RLC, CEF; Project administration: RLC, CEF; Funding acquisition: RLC, CEF.

